# Resources for genome editing in livestock: Cas9-expressing chickens and pigs

**DOI:** 10.1101/2020.04.01.019679

**Authors:** Denise Bartsch, Hicham Sid, Beate Rieblinger, Romina Hellmich, Antonina Schlickenrieder, Kamila Lengyel, Krzysztof Flisikowski, Tatiana Flisikowska, Nina Simm, Alessandro Grodziecki, Carolin Perleberg, Christian Kupatt, Eckhard Wolf, Barbara Kessler, Lutz Kettler, Harald Luksch, Ibrahim T. Hagag, Daniel Wise, Jim Kaufman, Benedikt B. Kaufer, Angelika Schnieke, Benjamin Schusser

## Abstract

Genetically modified animals continue to provide important insights in biomedical sciences. Research has focused mostly on genetically modified mice so far, but other species like pigs resemble more closely the human physiology. In addition, cross-species comparisons with phylogenetically distant species such as chickens provide powerful insights into fundamental biological and biomedical processes. One of the most versatile genetic methods applicable across species is CRISPR/Cas9. Here, we report for the first time the generation of Cas9 transgenic chickens and pigs that allow *in vivo* genome editing in these two important agricultural species. We demonstrated that Cas9 is constitutively expressed in all organs of both species and that the animals are healthy and fertile. In addition, we confirmed the functionality of Cas9 for a number of different target genes and for a variety of cell types. Taken together, these transgenic animal species expressing Cas9 provide an unprecedented tool for agricultural and biomedical research, and will facilitate organ specific reverse genetics as well as cross-species comparisons.

**Significance statement:** Genome engineering of animals is crucial for translational medicine and the study of genetic traits. Here, we generated transgenic chickens and pigs that ubiquitously express the Cas9 endonuclease, providing the basis for *in vivo* genome editing. We demonstrated the functionality of this system by successful genome editing in chicken and porcine cells and tissues. These animals facilitate organ specific *in vivo* genome editing in both species without laborious germ line modifications, which will reduce the number of animals needed for genetic studies. They also provide a new tool for functional genomics, developmental biology and numerous other applications in biomedical and agricultural science.

## Introduction

Chickens and pigs are the most important livestock species worldwide. They are not only important sources of food, but also valuable models for evolutionary biology and biomedical science. Pigs share a high anatomical and physiological similarity with humans, and are an important species for translational biomedical research e.g. in the areas of cancer, diabetes, neurodegenerative and cardiovascular diseases (1–3). In contrast, chickens are phylogenetically distant vertebrates from humans, but they were instrumental in the field of developmental biology due to the easy access to the embryonated egg. They are used to study neurological and cardiovascular functions (4–6) and provided key findings in B cell development and graft versus host responses (7–9).

Modelling human diseases in animals helps elucidating disease pathways and enables the development of new therapies. Although mice are an intensively studied vertebrate model (10), they are often not optimal for modelling particular human diseases. For example, mouse models for familiar adenomatous polyposis (FAP) poorly reflected the human pathophenotype since they develop polyps in the small intestine instead of the large intestine as observed in human patients (11) or in a porcine FAP model (12). Genetically modified livestock species hold great promise not only for biomedicine but also for agriculture. This has been demonstrated through new approaches for disease control, such as the generation of genome-edited pigs and chickens resistant to Porcine Reproductive and Respiratory Syndrome (PRRS) and Avian Leucosis Virus (ALV) infection, respectively (13–15).

Although the chicken was the first livestock animal with a full length genome available (16), precise manipulation of the chicken genome has been difficult and only became possible due to recent improvements in the cultivation of primordial germ cells (PGCs) (17, 18). The generation of pigs with precise germline-modifications required gene targeting in somatic cells followed by somatic cell nuclear transfer, which are both challenging, inefficient and time-consuming steps. Efficiency could be improved with the advent of the synthetic endonucleases such as Cas9. Cas9 (CRISPR associated protein 9) is a RNA-guided DNA endonuclease enzyme, associated with CRISPR (Clustered Regularly Interspaced Short Palindromic Repeats) (19) that has been extensively used for precise genome editing in different species including bacteria, plants and several animal species (20–24).

However, simultaneous delivery of Cas9 and sgRNAs to a specific organ *in vivo* has been taxing as the size of the *Cas9* gene challenges the packaging capacity of both adeno-associated virus (AAV) and lentivirus vectors (25, 26). To circumvent this drawback, Cas9 transgenic mice have been generated, requiring delivery of only the respective sgRNAs (27). The option to carry out *in vivo* genome editing provides not only the advantage that a gene of interest can be directly inactivated in the organ of choice but also bypasses the need for the generation of germline-modified animals, an important consideration when it comes to livestock species. We therefore decided to generate both Cas9 transgenic pigs and chickens that ubiquitously express Cas9 endonuclease. These provide an innovative and efficient model for *in vivo* genome editing as well as for *ex vivo* experiments, e.g. editing of primary cells or organoids.

## Results

### Generation of Cas9 transgenic animals

In this study, we generated transgenic pigs and chickens that ubiquitously express *Streptococcus pyogenes* Cas9 (SpCas9).

### Cas9 transgenic pigs

were generated by targeted placement of SpCas9 at the *ROSA26* locus (Figure 1a), which was previously shown to support abundant ubiquitous transgene expression in pigs (28, 29). 5% of G418-resistant cell clones showed correct gene targeting and were used for somatic cell nuclear transfer resulting in two liveborn piglets (#41, #42) (Figure 1c). Correct targeting was shown for both animals by long-range PCR across the 5’and 3’ junctions of the targeted allele and DNA sequence analysis. Monoallelic targeting was shown by PCR across the non-targeted *ROSA26* wildtype allele (supplementary data S1). Pig #42 served as a founder animal of a SpCas9 transgenic herd.

**Figure 1:**
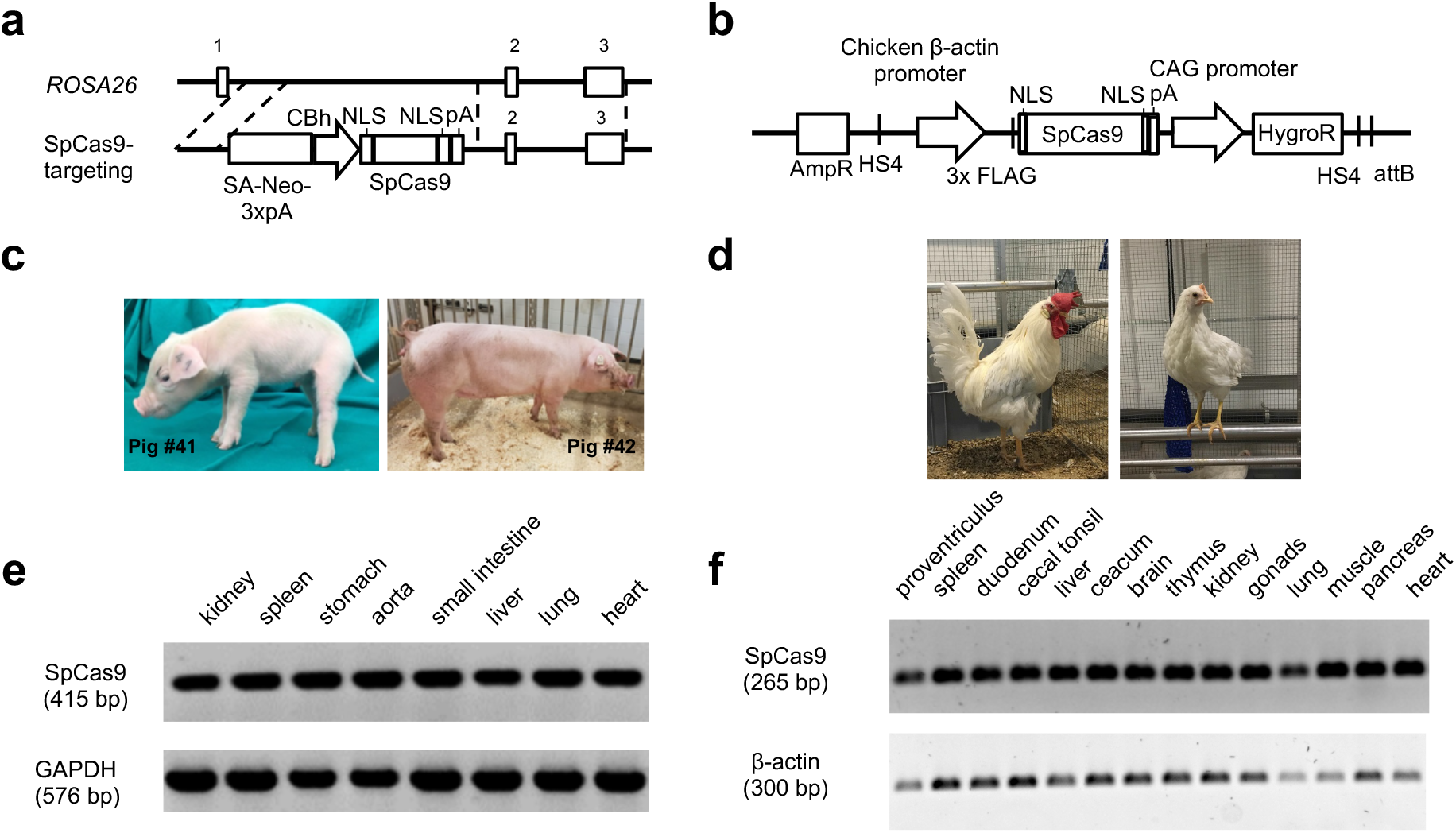
Generation and expression analyses of SpCas9 transgenic pigs and chicken. a) Structure of the ROSA26-SpCas9 targeting vector and targeting strategy to introduce *SpCas9* gene into the porcine *ROSA26* locus. Exons are indicated by numbered boxes and regions of homology by dotted lines. b) Expression vector used for the generation of SpCas9-expressing PGCs. c) SpCas9 transgenic founder pigs. d) SpCas9 transgenic rooster and hen. e) RT-PCR analyses of SpCas9 transgenic pig organs. SpCas9 expression is shown by a 415 bp PCR product and porcine GAPDH (576 bp) serves as a control. f) RT-PCR analyses of SpCas9 transgenic chicken organs. SpCas9 expression in all tissues analysed is shown by a 265 bp PCR product and β-actin (300 bp) serves as a control.

### Cas9 transgenic chickens

were generated by phiC31 integrase-mediated integration of SpCas9 into a chicken endogenous pseudo attP site, which has previously been reported to increase the integration frequency in PGCs (30) (Figure 1b). The PGCs carried the enhanced green fluorescent protein (EGFP) transgene. Cas9-EGFP-PGCs clones were screened for SpCas9 expression by immunofluorescence and flow cytometry (supplementary data S2a). Functionality was tested prior to generating germ line chimeras by transient transfection of Cas9-EGFP-PGCs with an expression vector for a sgRNA targeting the EGFP gene. Analysis by flow cytometry showed that 17% of SpCas9-expressing cells lost EGFP fluorescence, while no reduction was observed for the MOCK control (supplementary data S2b). The injection of modified PGCs (PGC clone 4) into 65 h old embryos resulted in 19 chimeric germline roosters. Upon sexual maturity, chimeric germline roosters were mated with wild type hens to obtain fully transgenic SpCas9 chickens (Figure 1d).

The transgene copy number was determined in both species via droplet digital (dd) PCR and revealed a single copy of SpCas9 in pigs and in chickens (supplementary data S3a, b). Both Cas9 transgenic chicken and pigs developed normally, showed no obvious abnormalities (i.e. in weight gain, supplementary data S3c) and were fertile.

### Ubiquitous expression of SpCas9 in transgenic animals

We analysed SpCas9 mRNA expression in different tissues from chickens (proventriculus, spleen, duodenum, cecal tonsil, caecum, liver, kidney, lung, pancreas, heart, brain, thymus, muscle and gonads) and pigs (stomach, spleen, small intestine, aorta, liver, kidney, lung and heart). Analysis of the house keeping genes glyceraldehyde 3-phosphate dehydrogenase (*GAPDH*) for pig samples and β-actin for chicken samples served as expression controls. The Reverse-Transcriptase-(RT-) PCR showed consistent SpCas9 expression in all samples of both species (Figure 1e, f).

### Functionality of SpCas9

To examine the functionality of SpCas9, we isolated different primary cell types from Cas9-expressing animals and transfected those with sgRNAs directed against EGFP, β2-microglobulin (*B2M),* a component necessary for the assembly of the MHC class I complex or against the C-X-C chemokine receptor type 4 (*CXCR4*) for the avian experiments. Targets in porcine cells were the endogenous *B2M* gene and the α-1,3-galactoyltransferase (*GGTA1*) gene responsible for the presentation of alpha Gal epitopes on the cell surface of most mammals, which play a crucial role in hyperacute rejection in pig-to-primate xenotransplantation (31, 32). Successful inactivation of these target genes can be assessed by flow cytometry.

### Cas9 transgenic pigs

Porcine ear fibroblasts (PEFs) were isolated from the founder pigs (#41 and #42), and transiently transfected with a vector containing a sgRNA against *B2M* (β2m-sgRNA).

Flow cytometry analyses revealed that 28.1% of cells from animal #41 lost expression of β2m and 20.6% of cells for pig #42 (Figure 2a).

Next, a SpCas9 transgenic offspring was sacrificed, porcine adipose derived mesenchymal stem cells (PADMSCs), porcine aorta endothelial cells (PAECs) and porcine kidney fibroblasts (PKFs) were isolated and transiently transfected with a vector containing a sgRNA against *B2M*. Flow cytometry analyses revealed that 63.1% of cells were negative for β2m expression in PADMSCs, 35.8% in PAECs and 16.2% in PKFs (Figure 2b), confirming the functionality of SpCas9 in those cells. Differences in editing efficiencies most likely correspond to different transfection efficiencies in these cell types.

**Figure 2:**
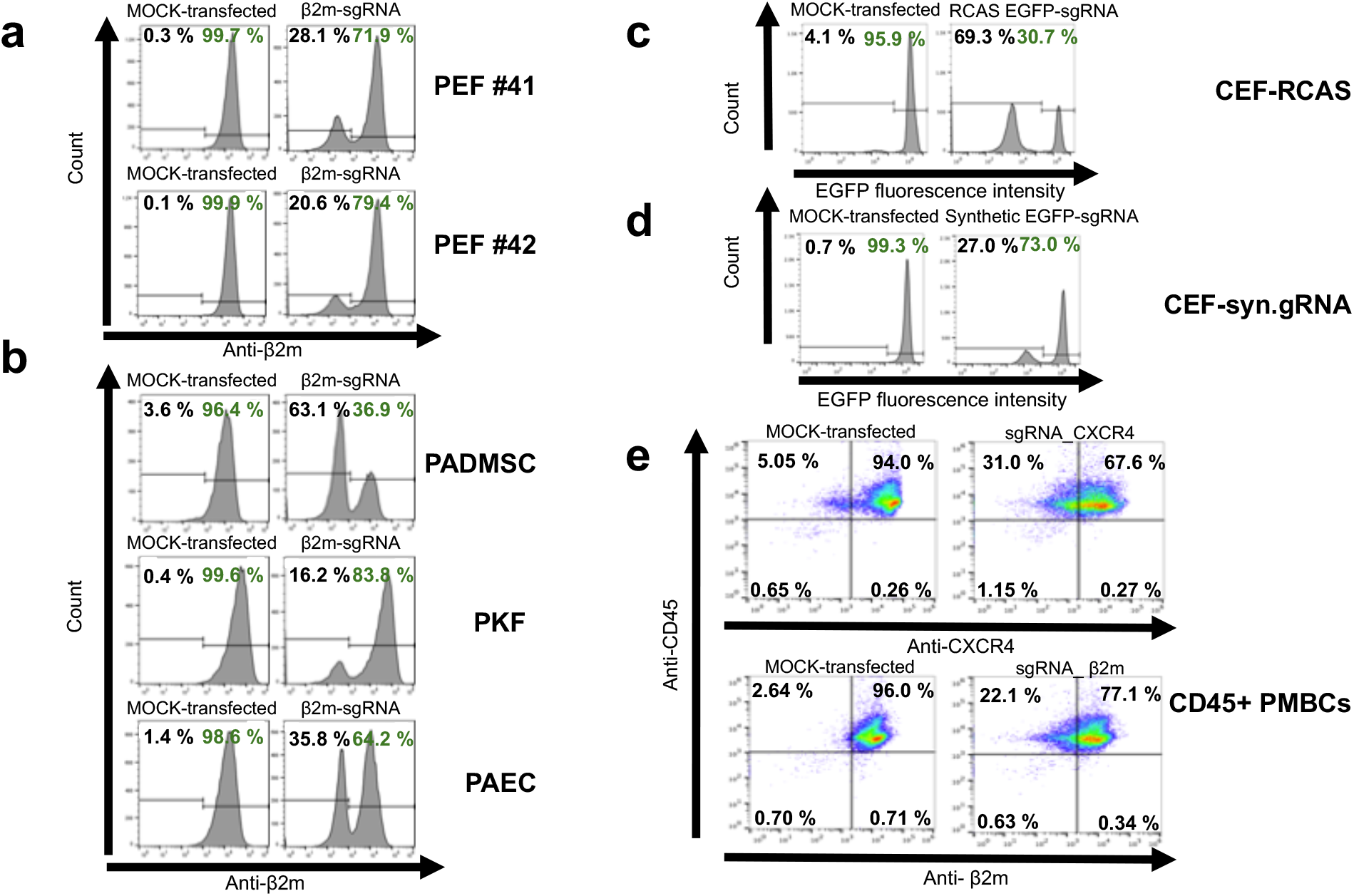
SpCas9 functionality in primary cells derived from SpCas9 transgenic pigs and chickens. a) PEFs derived from ear tissue of SpCas9 founder pigs #41 and #42 transfected with a construct carrying a sgRNA against *B2M* or *GGTA1* (MOCK-transfected control). b) PADMSCs, PKFs, PAECs derived from an SpCas9 transgenic piglet and transfected with a construct carrying a sgRNA against *B2M* or *GGTA1* (MOCK-transfected control). c) CEFs derived from a SpCas9-expressing chicken and transfected with RCASBP-(A)-EGFP-sgRNA or RCASBP-(A)-INF- sgRNA (MOCK-transfected control) d) CEFs transfected with a synthetic sgRNA against EGFP or *B2M* (MOCK-transfected control). e) Splenic CD45+ PMBCs derived from a SpCas9-expressing chicken and transfected with chemically modified gRNA against *B2M* or *CXCR4*.

Finally, to assess *ex vivo* functionality of SpCas9 in a more complex tissue, porcine colonic organoids were isolated from normal colonic mucosa of a SpCas9-expressing pig and transiently transfected with a vector carrying a sgRNA directed against the *GGTA1* gene. Tracking of Indels by DEcomposition (TIDE) analysis of a sequenced PCR fragment across the sgRNA target site revealed a total target cleavage efficiency of 12.8% in porcine colonic organoids (Figure 3a).

**Figure 3:**
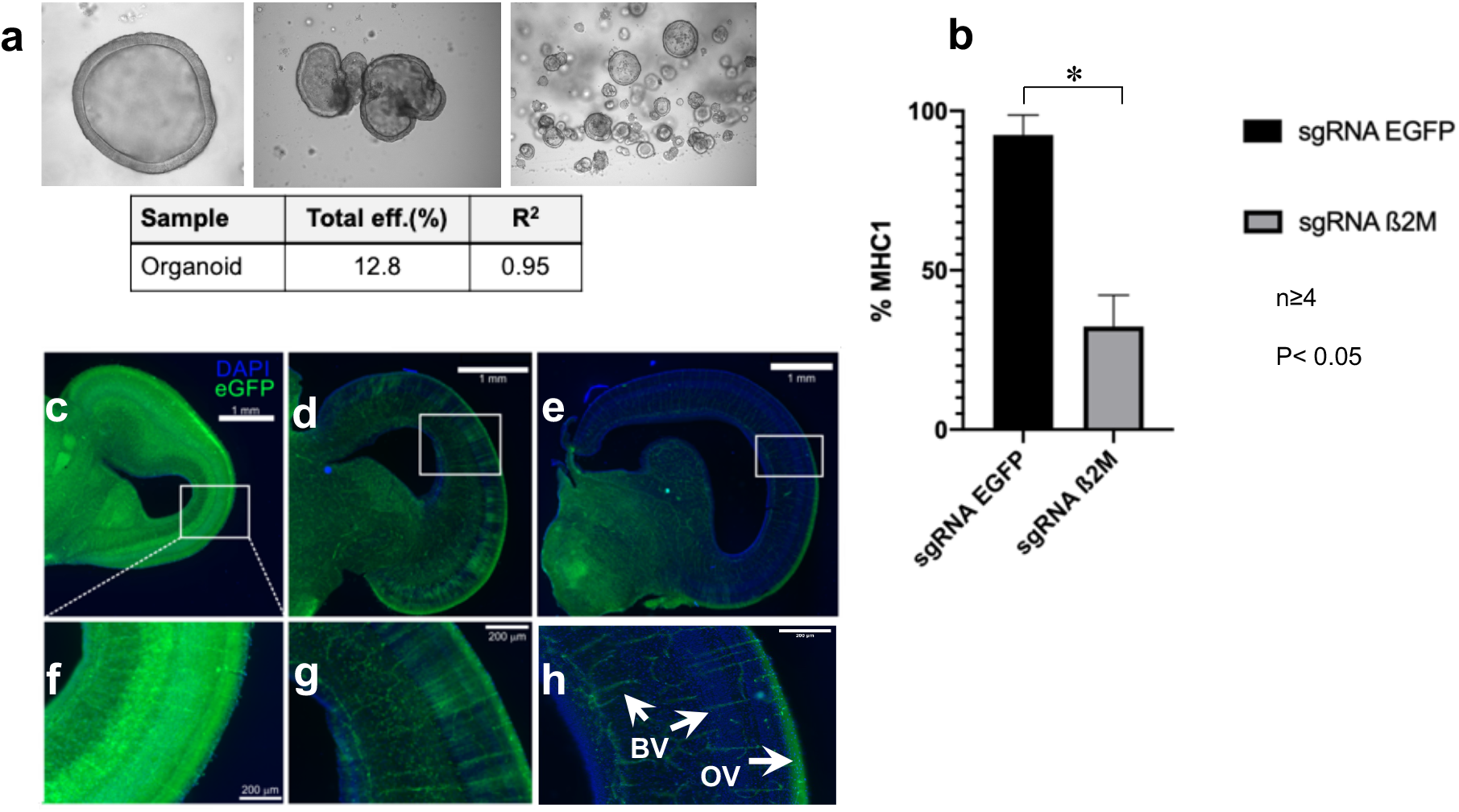
*Ex vivo* genome editing in porcine colonic organoids and *in ovo* genome editing in chickens. a) *Ex vivo* genome editing in porcine colonic organoids derived from an SpCas9 transgenic pig and transfected with a construct carrying a sgRNA against *GGTA1*. Top: Pictures of porcine colonic organoids, bottom: Genome editing efficiencies in porcine colonic organoids analysed by TIDE analysis. b) *In ovo* genome editing of *B2M* by RCASBP-(A) mediated gRNA delivery. At ED3, embryonated eggs were injected with RCASBP-(A)- β2m-sgRNA infected DF- 1 cells or RCASBP-(A)-EGFP-sgRNA infected DF-1 cells (MOCK control). At ED18, bursal B cells were analysed for β2m expression by flow cytometry (n≥4). P-value <0.05. c) *In vivo* genome editing via i*n ovo* electroporation in chickens. At ED2, midbrain vesicles of EGFP positive embryos were electroporated with pBlueScript II SK (+) vector containing a sgRNA against EGFP or β2m and embryos collected at ED 12. DAPI treated midbrain sections were analyzed for EGFP expression by fluorescence microscopy. Coronal section of midbrain optic tectum of MOCK β2m- sgRNA transfected embryo with clear EGFP signal in every cell (neurons, glia, and blood vessels). d, e) Frontal sections of optic tectum of EGFP-sgRNA transfected embryos showing a prominent reduction of EGFP fluorescence. f,g,h) Magnified sections from c,d,e illustrating tectal layering and loss of EGFP in g and h. Blood vessels (BV) and optical input fibers (OF) still expressing EGFP marked with white arrows.

### Cas9 transgenic chicken

Primary chicken embryonic fibroblasts (CEFs) were derived from an EGFP/Cas9 transgenic embryo. SgRNA against EGFP was delivered by the retroviral RCASBP-(A) (Replication Competent ALV LTR with a Splice acceptor) vector, which is a derivative of the avian Rous sarcoma virus (33, 34). Six days post transduction, 69.3% of cells were negative for EGFP expression as measured by flow cytometry (Figure 2c), while transfection of a synthetic unmodified sgRNA against EGFP resulted in 27.0% loss of EGFP expression after 48 h (Figure 2d). These results confirmed that viral transduction is more efficient in primary cells compared to transfection.

Next, genome editing was demonstrated in primary CD45+ peripheral blood mononuclear cells (PBMCs) derived from a spleen of a Cas9-expressing chicken. Cells were electroporated with two 2’-O-methyl phosphorothioate linkage-modified synthetic sgRNAs directed against *CXCR4* or *B2M* resulting in a reduction of 31% in CXCR4 expression and 22.1% in β2m expression in a CD45+ co-stained cell population (Figure 2e).

### Genome editing in embryonated chicken eggs

To examine the efficiency of genome editing *in ovo*, 3 days old embryonated eggs were injected in the allantois cavity with a DF-1 cell line producing the infectious RCASBP-(A) vector for expression of the sgRNA against *B2M*. The β2m expression on bursa-derived B cells was then analysed by flow cytometry and showed significant reduction of β2m on the cell surface compared to MOCK-transduced controls (p<0.05) (Figure 3b). However, high variability in the number of β2m expressing cells was observed, ranging between 30 to 80%.

Moreover, we could show *in vivo* functionality of SpCas9 in *in ovo*-electroporated chicken embryos with vectors containing a sgRNA against *B2M* or EGFP. Fluorescence microscopy imaging clearly showed a loss of EGFP in Cas9 positive brains (Figure 3c). In contrast, no EGFP loss was observed in Cas9/EGFP embryos electroporated with sgRNA against *B2M*. TIDE analysis for EGFP targeting revealed a total cleavage efficiency of 11.5% (embryo #5) and 6.8% (embryo #7) (Supplementary data S4) and for *B2M* 13.8 % (embryo #6), 11.8 % (embryo #12) and 11.1 % (embryo #22) (Supplementary data S5).

## Discussion

The generation of genetically modified livestock for basic science, evolutionary biology, agriculture and translational medicine has been a laborious, inefficient and time-consuming procedure. In pigs this relied on gene targeting in somatic cells followed by somatic cell nuclear transfer, while genetically modified chickens were usually generated by injecting genetically modified primordial germ cells (PGCs) into the avian embryos. The resulting chimeric birds had to be bred further to obtain non-chimeric founder animals. The CRISPR/Cas9 technology has now provided the tools to improve efficiency of genome editing so that, for some experimental setups, generation of germline modified animals is no longer required, as genome editing can be carried out directly *in vivo* in specific organ or cells. To simplify *in vivo* genome editing in livestock, we have generated pigs and chickens that ubiquitously express SpCas9 in all organs.

The SpCas9-expressing pigs were generated by targeted integration of SpCas9 into the *ROSA26* locus. This locus is known to be a ‘safe harbour’ for transgene expression without interrupting the function of essential endogenous genes (35). Moreover, it is readily targeted, supports abundant ubiquitous expression, and is dispensable for normal physiology and development (28, 29). Unlike previous work, where a Cre-inducible Cas9-expressing line has been generated (36), our pig line expresses a single copy of SpCas9 ubiquitously under the control of the CBh promoter. This eliminates the need for two consecutive recombination events to occur and should increase editing efficiency. For example, only 0.1% of cells expressed both Cre-recombinase and the inducible Cas9 after *in vivo* lentiviral-mediated delivery of Cre (36). Ubiquitously Cas9-expressing pigs are healthy and fertile and could be bred to homozygosity, which agrees with findings made in *ROSA26*-Cas9-expressing mice (27). Obtaining a genetic modification in the generated Cas9-expressing animals requires the delivery of the small sgRNA that could be efficiently packaged into AAVs and lentiviruses. Moreover, AAVs provide a certain degree of tissue specificity by local delivery of appropriate serotypes (37) allowing tissue specific genome editing within these animals.

Importantly, we also report the generation of the first Cas9 transgenic chickens. These were obtained by PhiC31-mediated transgene integration into an endogenous pseudo attP site (38). We took advantage of the ability to culture and genetically modify EGFP-expressing PGCs that were used to colonize the gonads and transmit Cas9 and EGFP transgenes to the offspring. The generated chickens, which express SpCas9 from only a single copy at a random position, are healthy and fertile. Initially, we aimed for high Cas9 expression to ensure efficient genome editing. However, if cell clones with more than a single copy of SpCas9 were used to generate Cas9 transgenic birds, lethality of the embryo was observed between ED18 and ED21 (data not shown). This may have been due to insertional mutagenesis or more likely due to a potential toxic effect of high Cas9 expression as observed for Cre-recombinase expressing mice (39).

The functionality of SpCas9 was confirmed in a variety of different cell types derived from SpCas9-expressing pigs and chickens. Besides showing Cas9-mediated genome editing in a series of porcine primary cells, we demonstrated *ex vivo* functionality of SpCas9 in porcine colonic organoids. Our group has an interest in modelling human cancers in pigs such as colorectal cancer (12). As previously shown in mice (40), modification of colonic organoids from SpCas9-expressing pigs enables *ex vivo* local inactivation of single or multiple tumor-relevant genes and the analysis of oncogenic transformation prior to subsequent autologous re-transplantation. This procedure will eliminate the need to generate germline modified pigs and will allow closer simulation of human oncogenesis.

The generation of SpCas9-expressing chickens allows efficient genetic modification of primary chicken cells without the need to deliver the Cas9 endonuclease. We showed the possibility of genome editing in CEFs via RCAS-delivered sgRNAs and via synthetic sgRNAs (Figure 2c, d). To our knowledge, this is the first study that reports successful genome editing in chicken cells using synthetic guides. Although both approaches were proven to be successful, transfection with synthetic sgRNA has the advantage of higher biosafety while more cells were transduced using the viral vector.

The easy access to the chicken embryo enables the study of mechanisms governing evolution and development (41). It also provides the possibility of *in ovo* delivery of plasmid constructs (42). Here we showed *in vivo* genome editing in chicken embryos via *in ovo*-electroporation of brain tissue with plasmids expressing sgRNAs against *B2M* and EGFP.

As an alternative delivery system the retroviral RCAS vector was tested for *in ovo* genome editing of *B2M* in B cells. Although RCAS-mediated genome editing was detected in all transduced embryos, the efficiency was highly variable, presumably due to different transduction efficiencies in individual embryos.

Finally we demonstrated the possibility of genetic modification of cells that are usually hard to transfect such as lymphocytes (43). We used chemically modified gRNAs as they were shown to increase the stability of gRNAs in primary cell culture (44, 45), and were able to efficiently inactivate the *CXCR4* and *B2M*genes in splenic CD45 positive cells derived from Cas9-expressing chickens.

In conclusion, we generated a versatile new tool and showed efficient genome editing of four different target genes in a number of cell types and in the developing embryo, confirming SpCas9 functionality in both pig and chicken. We are confident that the SpCas9 transgenic pigs and chickens provide a powerful platform for *in vivo* genome editing in two important livestock species which also represent important animal models for developmental and biomedical research.

The generated Cas9-expressing animals provide an improved and efficient platform for genome editing, eliminate the need for Cas9 delivery *in vivo* and minimize the time, costs and number of animals needed for the generation of genome-edited animals.

## Methods

### Generation of SpCas9- and sgRNA constructs

For the generation of SpCas9 expression constructs, we used the sequence of a human codon optimized *Streptococcus pyogenes* Cas9 (SpCas9) obtained from pX330-U6-Chimeric_BB-CBh-SpCas9 from Feng Zhang (Addgene plasmid # 42230; http:/n2t.net/addgene:4223O;RRID:Addgene_42230). Oligonucleotides were synthesized by Eurofins Genomics, Germany.

For stable integration in chicken PGCs, the SpCas9 construct contained an attB site to ensure the insertion of the transgene using phiC31 integrase (38). SpCas9 was amplified by PCR and the PCR product was assembled with a −8.7 kb backbone construct using NEBuilder® HiFi DNA Assembly Cloning Kit (New England Biolabs, USA) according to manufacturer’s instructions. The SpCas9 expression cassette contained a 1.3 kb chicken β-actin promotor, followed by SV40 NLS, a 4.1 kb *SpCas9* gene, a nucleoplasmin NLS and a SV40 poly A. The generated construct contained a CAGGS-hygromycin resistance cassette and the *SpCas9* gene was flanked by duplicated copies of the core 300-bp HS4 insulator from the chicken β-globin gene to ensure proper transgene expression (Figure 1b).

For precise placement of SpCas9 at the porcine *ROSA26* locus, we generated a ROSA26-SpCas9 targeting vector by cloning a 5.3 kb fragment containing the chicken β-actin hybrid intron (CBh) promoter-driven SpCas9 construct between both homologous arms of a previously generated ROSA26-promoter trap vector (28). The generated ROSA26-SpCas9 promoter trap vector consisted of: a 2.2 kb 5’ homology arm corresponding to a region of porcine *ROSA26* intron 1; a 1.6 kb fragment composed of splice acceptor, Kozak sequence, promoterless neomycin resistance gene and a triple-poly-A signal; a 5.3 kb SpCas9 expression cassette; and a 4.7 kb 3’ homology arm corresponding to a region of porcine *ROSA26* intron1-3. The SpCas9 cassette contained a 0.8 kb CBh promoter, followed by a SV40 NLS, a 4.1 kb *SpCas9* gene, a nucleoplasmin NLS and a 0.2 kb BGHpA (Figure 1a). The CBh promotor was composed as described earlier (46).

SgRNA constructs directed against EGFP and chicken *B2M* were generated by cloning the respective gRNA oligonucleotides (EGFP: 5’-CCGGCAAGCTGCCCGTGCCC-3’; chicken B2M: 5’-GAACGTCCTCAACTGCTTCG- 3’) with additional BbsI overhangs into a BbsI-digested pBluescript II KS (+) vector carrying a 0.2 kb U6 promotor and a sgRNA scaffold sequence. SgRNA constructs directed against porcine *GGTA1* and *B2M* were generated by cloning the respective gRNA oligonucleotides (GGTA1 exon 7: 5’- GTCGTGACCATAACCAGA- 3’; porcine B2M exon 1: 5’- TAGCGATGGCTCCCCTCG- 3’), with additional BbsI overhangs into a BbsI-digested vector carrying an 0.2 kb U6 promotor, a sgRNA scaffold sequence and additionally a 1.6 kb puromycin resistance cassette composed of a 0.8 kb CBh promoter, a 0.6 kb puromycin resistance gene and a 0.2 kb BGHpA. Oligonucleotides were synthesized by Eurofins Genomics, Germany. The generated vectors consisted of: a U6 promoter followed by the gRNA sequence and a sgRNA scaffold sequence; and a puromycin resistance cassette for sgRNA constructs directed against porcine *GGTA1* and *B2M* (Figure 2a, b).

In chickens, sgRNAs were also delivered by using the RCAS system. The sequences coding for sgRNAs and a human U6 promoter were introduced into the ClaI restriction site of the RCASBP-(A) vector as previously described (47, 48) and correct insertion was verified by sequencing (Eurofins Genomics, Germany). Synthetic gRNAs were synthesized by SYNTHEGO (Synthego, USA) and prepared according to manufacturer’s instructions.

### Generation of SpCas9-expressing cells

Gene targeting in porcine cells was performed in porcine kidney fibroblasts that were isolated and cultured by standard methods (49). Cells were transfected with 4-6 μg linearized targeting vector DNA (ROSA26-SpCas9) using Lipofectamine 2000 transfection reagent (Thermo Fisher Scientific, USA), selected with 1400-1600 μg/mL G418 (Genaxxon Bioscience, Germany) for 7-10 days, and targeted clones identified by 5’- and 3’-junction PCR and DNA sequence analysis.

The generation of transgenic chickens required genome editing of PGCs which colonize the gonad, creating chimeric germline animals. In this study, EGFP-PGCs were derived from the germinal crescent as previously described (50). They were maintained in culture and stably transfected with a Cas9 construct which was mediated by phiC31 integrase as previously described (51). Briefly, 5 × 10^6^ of EGFP-PGCs were resuspended in a total volume of 100 μl Nucleofector Solution V (Lonza, Switzerland) premixed with 10 μg of Cas9 plasmid and the same amount of phiC31 integrase. Electroporation was conducted by ECM 830 Square Wave Electroporation System (BTX) using eight square wave pulses (350 V, 100 μsec). After transfection, cells were plated on 48 well plates and selected with 50 μg/mL hygromycin.

### Animals

Permission for the generation of transgenic Cas9-expressing chickens and pigs was issued by the government of Upper Bavaria, Germany (ROB-55.2-2532.Vet_02-17-101 and ROB-55.2-2532.Vet_02-18-33 for transgenic chickens and pigs, respectively). Experiments were performed according to the German Welfare Act and European Union Normative for Care and Use of Experimental Animals. All animals received standard diet and water ad libitum.

Transgenic pigs were generated as previously described (52). Briefly, donor cells were arrested at G0/G1 phase by serum deprivation. Oocytes, isolated from prepubertal gilts, were maturated *in vitro*, enucleated and single donor cells inserted into the perivitelline space. Cells were fused and oocytes activation was induced via electric pulse. Reconstructed embryos were then transferred into the oviducts of hormonally synchronised recipient sows.

Transgenic chickens were generated as previously described (53). Briefly, 3000 Cas9-PGCs were injected into the vasculature of 65 h old embryos (Lohmann selected White Leghorn line (LSL), Lohmann-Tierzucht GmbH, Germany), subsequently transferred into a turkey surrogate shell and incubated until hatch of chimeric roosters. Upon sexual maturity, sperm was collected for DNA isolation and genotyping. After breeding with wild type hens, fully Cas9 transgenic chickens were obtained.

### Derivation and functional assays in SpCas9-expressing cells

The functionality was tested in SpCas9-expressing cells and various cell types derived from the SpCas9-expressing animals.

Porcine ear fibroblasts (PEFs), porcine adipose derived mesenchymal stem cells (PADMSCs), porcine aortic endothelial cells (PAECs) and porcine kidney fibroblasts (PKFs) were isolated according to standard protocols (49, 54, 55) from a SpCas9-expressing pig and cultured in Dulbecco’s Modified Eagle Medium (DMEM) (PEFs and PKFs) or Advanced DMEM (PADMSCs and PAECs) supplemented with 20% fetal bovine serum, 2 mM Ala-Gln, 1x MEM non-essential amino acid solution and 1 mM sodium pyruvate (Corning, USA) at 37 °C and 5% CO2. Cells were then transfected with 1 μg of sgRNA constructs against porcine *B2M* or *GGTA1* using Lipofectamine 2000 transfection reagent and 24 h post transfection, transiently transfected cells were selected using 1.5 μg/mL puromycin (InvivoGen) for 2 days. The remaining cell pool was analysed by flow cytometry (Figure 2a, b).

To test SpCas9-expressing PGCs before injection into the embryo, cells were transfected with sgRNA directed against EGFP and analysed by flow cytometry; 4×10^6^ cells were electroporated with 10 μg DNA of pBluescript II SK(+) carrying a human U6 promoter and EGFP-sgRNA using Nucleofector® V Kit (Lonza, Switzerland) as described above.

Chicken embryonic fibroblasts (CEFs) were isolated as previously described (56). Briefly, fertilized eggs were incubated at 37.8 °C. On day 10, −5 μl blood were carefully drawn for genotyping. The next day, CEFs were prepared from Cas9 positive embryos and resuspended in Iscove’s medium supplemented with 5% fetal bovine serum (FBS), 2% chicken serum and 1% Penicillin-Streptomycin. CEFs were incubated at 40°C under 5% CO2 conditions and subsequently used for functional assay with RCASBP-(A)-sgRNA as well as synthetic sgRNA. Transfection with RCAS-sgRNAs was performed using ViaFect™ Transfection Reagent (Promega, USA) according to manufacturer’s instructions with slight modifications. Briefly, CEFs were derived from EGFP-Cas9 and EGFP embryos and 2.5×10^5^ cells each were seeded in 6 well plates. Twenty four hours later, transfection complexes were prepared in a ratio of 6:1 (transfection reagent:DNA) where a total amount of 500 ng DNA was used and incubated for 20 min at room temperature. Transfection with synthetic sgRNAs was conducted using Xfect^TM^ RNA Transfection Reagent Protocol-At-A-Glance (Takara, Japan) following the manufacturer’s instructions. Briefly, 100,000 EGFP-Cas9 CEF cells were seeded in a 24 well plate and transfected 24 h later with 25 pmol of synthetic EGFP-sgRNA premixed with Xfect RNA transfection polymer. The complex was removed 4 h later and replaced by 500 μl cell culture medium. Cells were analysed at different time points for INDELs and loss of EGFP expression.

Splenic CD45+ cells were derived from a Cas9-expressing animal and isolated as previously described (57). Briefly, the spleen was removed aseptically and homogenized by passing it through a cell strainer. The cells were isolated by Ficoll density gradient separation and directly used for electroporation with 25pmol of chemically modified gRNAs directed against *CXCR4* (CXCR4_1434: A*A*A*UUCAAUGAGUAUGCCAG) or *B2M* (β2m_1444:U*C*U*UGGUGCCCGCAGAGGCG) (Synthego, USA). For electroporation, the Human T Cell Nucleofector Kit (Lonza, Switzerland) was used according to manufacturer’s instructions. The cells were plated onto a 12 well plate with RPMI, 10% FBS, 1% Penicillin/Streptomycin and 1% Glutamax. The medium was replaced with fresh medium 6 hours after electroporation. 48h post electroporation, the cells were analysed by flow cytometry.

### *In ovo* functionality of SpCas9 in chicken bursal B cells

Transduction of embryonic B cells was carried out using the RCAS retroviral gene-transfer system that was used as previously described (34, 58–60) (Figure 3b). Briefly, DF-1 cells were transfected with RCASBP-(A)-β2m-sgRNA or MOCK RCASBP-(A)-EGFP-sgRNA using ViaFect^TM^ Transfection Reagent (Promega, USA) according to manufacturer’s instructions. The transfected cells were cultured for 10 days to ensure completed infection. 1×10^6^ cells in 100 μl DMEM per egg were injected into ED3 old embryo using a sterile 1 ml syringe. At ED10, blood was taken from the embryos and genotyping was performed in order to identify SpCas9 positive embryos. At ED18, heart tissue was collected for insertion and deletion (INDEL) analysis and RCAS detection by PCR with primers RCAS_B2M_F (5’- CGAAGCAGTTGAGGACGTTC- 3’) and RCAS_B2M_R (5’ -CATATTTGCATATACGATACAAGGC- 3’) resulting in a 245 bp amplicon. The bursa was dissected from the embryo to obtain B cells isolated by density gradient centrifugation. Since β2m is a necessary component for the assembly of the MHC I complex, inactivating β2m leads to the loss of MHC I at the cell surface. Thus, the cells were stained for β2m and analysed by flow cytometry.

### *Ex vivo* functionality of SpCas9 in porcine colonic organoids

*Ex vivo* functionality of SpCas9 was tested in porcine colonic organoids. Colonic organoids were isolated as follows: Colonic mucosa was incubated in dissociation buffer (PBS, 30 mM EDTA, and 10 mM DTT) for 20 min at 4°C, colonic crypts then mechanically separated under a microscope and embedded in Matrigel (Corning, USA). Organoids were cultivated in growth medium as previously described (61) with minor changes (50% mouse Wnt3a-conditioned medium, 50ng/mL mouse Wnt3a recombinant protein (Biolegend, USA), 15% mouse R-spondin1-conditioned medium, 1.3% porcine Noggin-conditioned medium, 2.5 μM CHIR99021 (Calbiochem/Merck, Germany), 100 μg/mL primocin (Invitrogen, USA).

Colonic organoids were transfected with 2 μg of sgRNA construct as previously described (62). Electroporation was performed using the BTX ECM 830 with following instrument settings: 180 V, 2 pulses of 5 ms length and 100 ms pulse interval. 24 hours post transfection, transiently transfected cells were selected using 1 μg/mL puromycin (InvivoGen) for 48 hours and cell pools were analysed for the presence of INDELs.

### *In vivo* functionality of SpCas9 in chicken central nervous system

Central nervous tissue (neurons and glial cells) was transfected using *in ovo* electroporation. Electroporation has previously been shown to be an effective method for targeted gene transfer in the developing chicken embryo (42, 63–66). Briefly, embryos of Cas9xWT and Cas9xEGFP at developmental stages HH 10 - 13 (after approximately 44h incubation) were transfected with pBlueScript II SK (+) vector containing a sgRNA directed against *B2M* or EGFP. To this aim, a stripe of adhesive tape was attached to the egg shell to prevent cracks, and about 2 ml albumen was suctioned with a syringe from the bottom of the egg. A window of about 1 to 2 cm^2^ was cut into the upper side to expose the blastodisc with the embryo. Blue light illumination of the blastodisc increased contrast under binocular microscope view.

The plasmid solutions were mixed 4:1 with Fast Green Dye resulting in 667 ng/μl of pBlueScript II SK (+) vector containing a sgRNA against *B2M* and 1255 ng/μl of pBlueScript II SK (+) vector containing a sgRNA against EGFP. A small volume of plasmid solution filling only the tip of a 100 μm glass micropipette was then injected into the mesencephalic vesicle (second brain vesicle). Next, a gold electrode (positive) was positioned at about 1 mm distance to the left of the embryo, and a tungsten micro-electrode (negative) was inserted into the second brain vesicle. For electroporation, 3 rectangular pluses at 2 Hz with 15 V each and 25 ms duration were applied (Grass S48 stimulator, Medical Instruments, USA with Genetrodes 45-0115, Harvard Apparatus Inc., MA, USA). Electrodes were removed and 1-2 ml of 4°C chicken ringer solution (150 mM NaCl, 5.4 mM KCl, 2.2 mM CaCl2, and 2.4 mM NaHCO3) dripped onto the embryo for liquid resupply and to reduce damage caused by heating. The eggshells were sealed with adhesive tape and the embryos were incubated until embryonic day 12, when the embryonic brains where dissected for genotyping, TIDE analysis and histological preparation of midbrain slices of EGFP positive embryos (Fig. 3c-h).

For histological analysis of EGFP-positive embryos, the ED12 midbrains including the optic tectum (OT) were fixed in 4% paraformaldehyde for at least 24 h, transferred into 30% sucrose solution for 4 h for cryo-protection, cryosectioned to 50 μm and mounted with DAPI-mounting medium (0.1mg DAPI in 100 ml n-propyl gallate). All images were taken with an epifluorescence microscope (Olympus BX63 with XM10 digital camera, Olympus, Germany) using the same exposure time and gain.

### INDEL detection

Cell pools were analysed for presence of INDELs by PCR spanning the sgRNA target site using primers presented in Table 1. The resulting amplicons were sequenced (Eurofins Genomics, Germany) and analysed with TIDE (Tracing of Indels by Decomposition) (https://tide.deskgen.com/) (67).

**Table 1:**
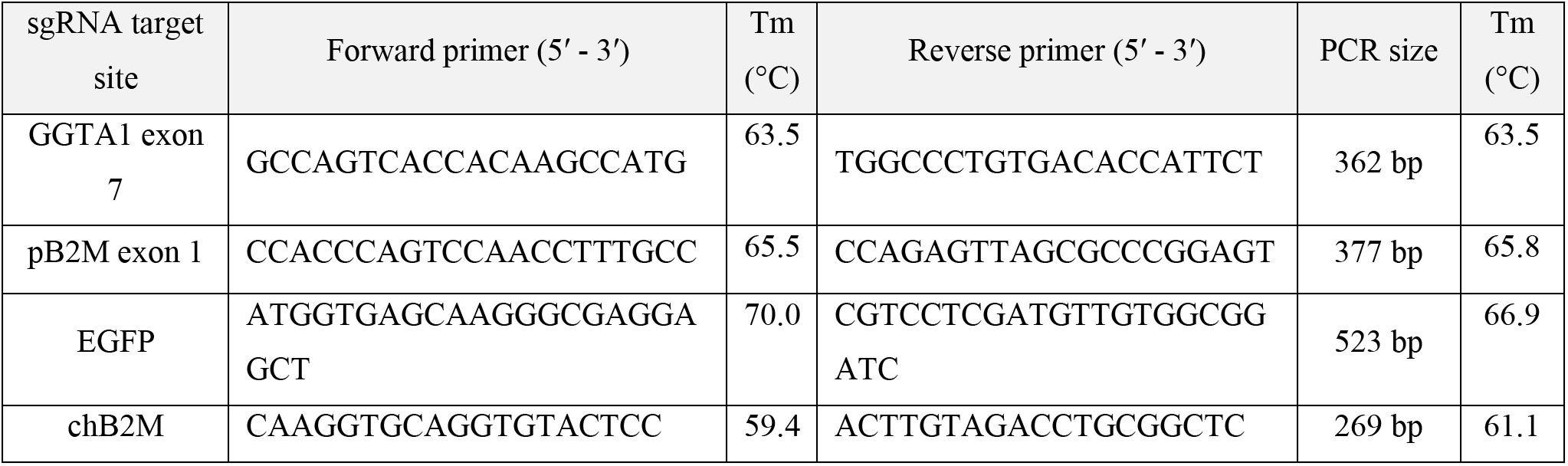
Primers used for INDEL detection.

### Genotyping PCRs

The following primers were used to identify correct placement of SpCas9 at the porcine *ROSA26* locus: 5’- junction PCR used primers ROSA26_5’F (5’- TATGGGCGGGATTCTTTTGC- 3’) and Neo_5’R (5’- AGCCCCTGATGCTCTTCGTC- 3’) to amplify a 3.1 kb fragment. 3’- junction PCR used primers Cas9_3’F (5’- GCAGATCAGCGAGTTCTCCA- 3’) and ROSA26_3’R (5’- CAGGTGGAAAGCTACCCTAGCC- 3’) to amplify a 5.6 kb fragment. The wild type *ROSA26* allele was amplified with primers ROSA26_5’F (sequence above) and ROSA26_I1R (5’- GTTTGCACAGGAAACCCAAG- 3’) producing a 3.1 kb fragment. PCRs were carried out using AccuStart Taq DNA polymerase HiFi (QuantaBio, USA) and thermal cycling conditions: 94°C, 1 minute followed by 40 cycles at 94°C for 20 sec; 60°C for 30 sec, 78°C for 1minute per kb, and final extension at 68°C for 5 minutes.

The following primers were designed for the specific detection of SpCas9 in transgenic chickens Cas9-593-For (5’-GAGAGAATGAAGCGGATCGAAGAG- 3’) and Cas-440-Rev (5’ -TTGCTGTAGAAGAAGTACTTGGCG- 3’). PCRs were carried out using FIREPol DNA Polymerase (Solis Biodyne, Estonia) and thermal cycling conditions: 95°C for 5 min followed by 40 cycles at 95°C for 30 seconds; 58 °C for 30 seconds, 72°C for 1 minute per kb, and final extension at 72°C for 5 minutes.

### Determination of the transgene copy number by droplet digital PCR (ddPCR)

ddPCR was performed as described previously (29). The transgene copy number was determined using a fluorescence-labelled probe for SpCas9 [(5’FAM- GCACGCCATTCTGCGGCGGC- BHQ3’); primers used: ddSpCas9_F (5’- AGTTCATCAAGCCCATCCTG- 3’) and ddSpCas9_R (5’- TCTTTTCCCGGTTGTCCTTC- 3’)] or hygromycin [(5’FAM -TCGTGCACGCGGATTTCGGCTCCAA- 3’); primers used: ddHygro_F (5’-CATATGGCGCGATTGCTGATC- 3’) and ddHygro_R (5’ -GTCAATGACCGCTGTTATGC- 3’)]. GAPDH [(5’HEX- TGTGATCAAGTCTGGTGCCC- BHQ3’); primers used: ddGAPDH_F (5’- CTCAACGACCACTTCGTCAA- 3’) and ddGAPDH_R (5’-CCCTGTTGCTGTAGCCAAAT- 3’)] or β-actin [(5’HEX-GTGGGTGGAGGAGGCTGAGC- BHQ3’); primers used: ddBeta_actin_F:)5’-CAGGATGCAGAAGGAGATCA- 3’) and ddBeta_actin_R: (5’-TCCACCACTAAGACAAAGCA- 3’)] were used as references. The target gene was quantified by using the QX100 system (Bio-Rad Laboratories).

### Transgene expression analysis

RNA from porcine samples was isolated using Sure Prep RNA/DNA/Protein Purification Kit (Fisher Scientific, USA) for cells, or the innuSPEED tissue RNA kit (Analytic Jena, Jena, Germany) for tissues and cDNA synthesized using the FastGene Scriptase (Nippon Genetics, Germany). SpCas9 expression was detected with primers pCas9_F1 (5’- GCAGATCAGCGAGTTCTCCA- 3’) pCas9_R1 (5’- GGGAGGGGCAAACAACAGAT- 3’) resulting in a 415 bp amplicon. Porcine GAPDH was detected with primers GAPDHF (5’- TTCCACGGCACAGTCAAGGC- 3’) and GAPDHR (5’- GCAGGTCAGGTCCACAAC- 3’) resulting in a 576 bp amplicon. PCR was performed using GoTaq G2 DNA Polymerase (Promega, USA) according to manufacturer’s instructions.

RNA from chicken samples was isolated by Reliaprep™ RNA Tissue Miniprep System according to manufacturer instructions (Promega, USA), followed by cDNA synthesis using GoScript Reverse transcription mix (Promega, USA). SpCas9 expression was detected with primers chCas9_F1 (5’- GAGAGAATGAAGCGGATCGAAGAG - 3’) chCas9_R2 (5’-CAGTTCCTGGTCCACGTACATATC- 3’) resulting in a 150 bp amplicon. β-actin was detected with primers Beta_actin_F (5’- TACCACAATGTACCCTGGC- 3’) and Beta_actin_R (5’- CTCGTCTTGTTTTATGCGC- 3’) resulting in a 300 bp amplicon. PCR was performed using FIREPol DNA Polymerase (Solis Biodyne, Estonia) according to manufacturer’s instructions.

### Immunofluorescence

Cytoplasmic detection of Cas9 in the generated PGC clones was done by FLAG-Tag staining. Briefly, a total number of 5×10^5^ cells were centrifuged, washed with PBS and incubated with 100μl fixation buffer (eBioscience, USA) for 10 min. Cells were washed with 2% BSA in PBS (FLUO-Buffer) followed by permeabilisation with 0.1% Triton X-100 in PBS for 10 min. Cells were incubated for 1h with a mouse anti-FLAG M2 antibodies at a dilution of 1:500 in FLUO-Buffer. Subsequently, cells were washed in PBS and incubated with an Alexa 568 secondary anti-mouse IgG for 1 h in dark. Cells were washed in PBS, resuspended in 20 μl mounting medium supplemented with DAPI and immunofluorescence detected under a fluorescence microscope (ApoTome, Zeiss, Germany).

### Flow cytometry

Extracellular staining was carried out to detect β2m and CXCR4. Briefly, 3×10^5^ - 1×10^6^ cells were used per sample and washed with 2% BSA in PBS (FLUO-Buffer). To determine the living cell population, cells were incubated with Fixable Viability Dye eFluor 780 (eBioscience, USA) at dilution of 1:1000. After washing with FLUO-Buffer, primary antibodies (concentration shown in Table 2) were applied for 20 min. Cells were washed in FLUO-Buffer to remove unbound antibodies and incubated with conjugated secondary antibodies for 20 min. Subsequently, cells were again washed and analysed using an Attune flow cytometer (Thermo Fisher Scientific, USA). To detect CD45+ cells, a direct FITC-labeled primary antibody was used and the cells were washed with FLUO-Buffer before analysis. Antibodies and concentrations used are shown in Table 2.

**Table 2:** Antibodies used for flow cytometry.

| Antibody | Company/Origin | concentration |
| --- | --- | --- |
| Anti chicken β2m F21-21(68) | Cell culture supernatant | 1:5 dilution |
| Anti pig-B2M-Biotin clone B2M-02 | Sigma-Aldrich, USA (SAB4700015) | 20 μg/ml |
| Streptavidin-PE | BD, USA (554061) | 2.5 μg/ml |
| CD45-FITC | Biozol, Germany (SBA-8270-02) | 5 μg/ml |
| CXCR4 | Bio-rad, USA (MCA6012GA) | 0.2 μg/ml |
| H+L-APC | Biozol, Germany (MBS-MBS4151013) | 0.625 μg/ml |

For intracellular staining cells were fixed and permeabilized with 0.1% Triton X-100 in PBS for 10 min. The subsequent staining process was carried out as described above. Detection of EGFP signal loss in different cells was examined by quantifying the loss of EGFP signal. For all flow cytometry analyses, an Attune flow cytometer (Thermo Fisher Scientific, USA) was used. Data was analysed with FlowJo 10.4.1 software (FlowJo, LLC 2006-2017, USA).

### Statistical analysis

Statistical analysis was carried out using SPSS24 statistics (version 24.0.0.0) software (IBM, USA). Normally distributed data (Shapiro-Wilk test*p* > 0.05) were analysed by student’s T-Test. The Mann-Whitney-U Test was applied for not normally distributed data. P-values < 0.05 were considered as significant. Graphs were constructed using GraphPad Prism (version 8.0.1 145) (GraphPad Software, USA).

## Authors contributions

Designed research: H. Sid, B. Schusser, A. Schnieke, B. Rieblinger, B.B. Kaufer, J. Kaufmann, H. Luksch.

Performed research and analysed data: D. Bartsch, H. Sid, B. Rieblinger, K. Lengyel, A. Grodziecki, N.Simm, C. Perleberg, A. Schlickenrieder, R. Hellmich, K. Flisikowski, T. Flisikowska, C. Kupatt, E. Wolf, B. Kessler, D.Wise, I.T. Hagag, L. Kettler, Contributed new reagents or analytic tools: B.B. Kaufer, D. Wise, J. Kaufman, Wrote the paper: D. Bartsch, H. Sid, B. Rieblinger, A. Schnieke, B. Schusser, B.B. Kaufer

