## Supporting information for "Resources for genome editing in livestock: Cas9-expressing chickens and pigs"

### Supporting information (SI)

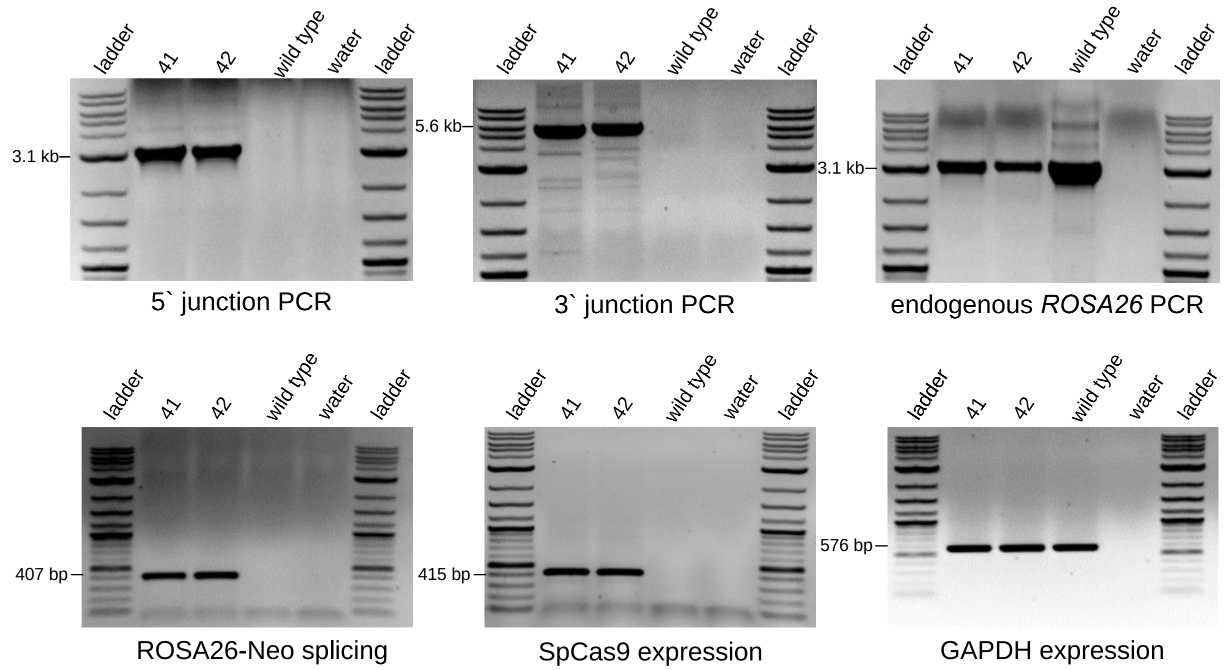


S1: Targeting PCR and SpCas9 expression analysis of piglets #41 and #42. a) Targeting PCRs performed to reveal correct targeting of 3.1 kb 5´junction, 5.6 kb 3´junction and monoallelic targeting by 3.1 kb endogenous PCR product. Wild type DNA was used as control. b) RT-PCR analysis of PEFs derived from SpCas9-expressing piglets #41 and #42. Correct splicing between *ROSA26* exon one and neomycin, SpCas9 and GAPDH expression reveals a 407 bp, 415 bp and 576 bp amplicon respectively. Wild type DNA was used as control.

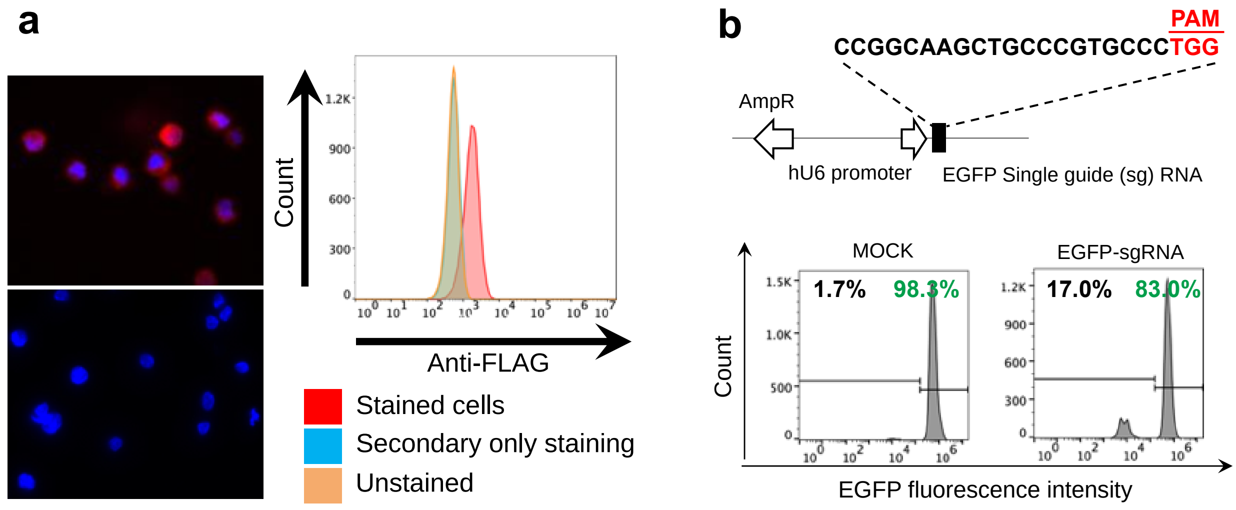


S2: Stable expression of SpCas9 und functionality in chicken PGCs. a) PGCs stained with anti-FLAG and analysed by immunofluorescence and flow cytometry. b) Flow cytometry analyses of PGCs electroporated with the pBlueScript II SK (+) vector containing a sgRNA against EGFP.

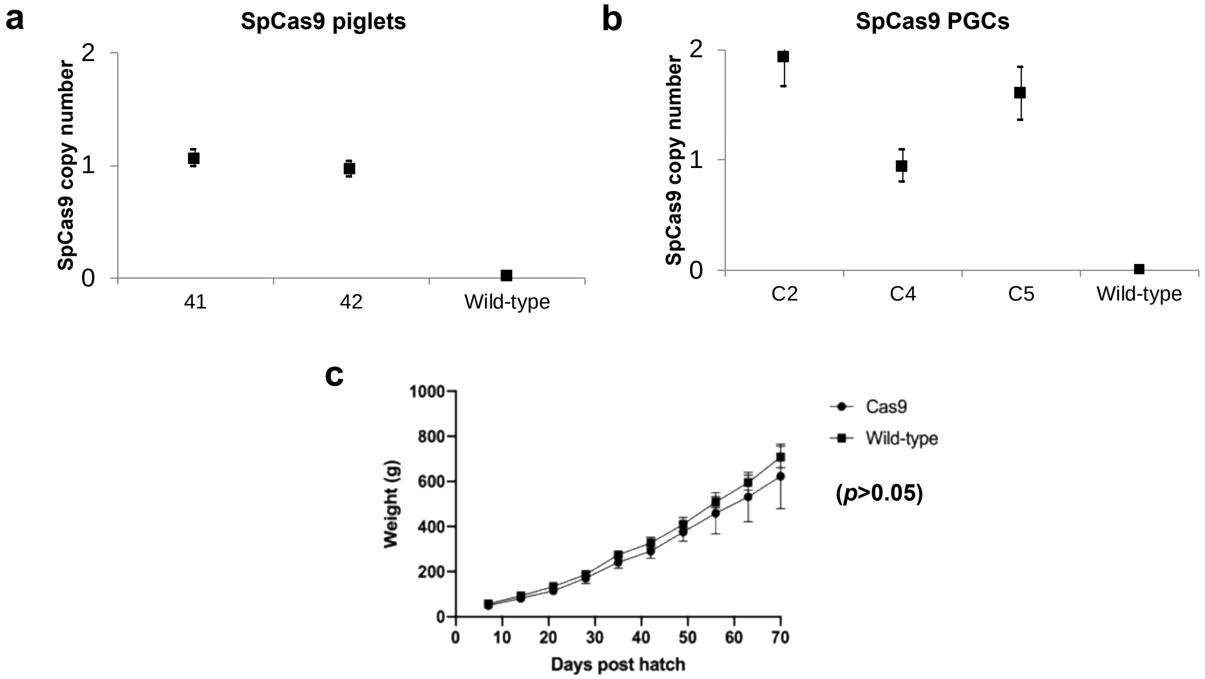


S3: SpCas9 copy number and development of SpCas9-expressing chicken and pigs a) Determination of SpCas9 copy number by droplet digital PCR in the SpCas9 transgenic founder pigs #41 and #42. Wild-type DNA was used as negative control. c) Determination of SpCas9 copy numbers by droplet digital PCR in chicken PGC clones C2, C4 and C5. Wild-type DNA was used as negative control. a) Weight comparison between SpCas9-expressing and wild-type chickens. Body weight was measured weekly for period of 10 weeks. SpCas9 transgenic chicken were healthy and showed no significant difference in weight gain compared to wild-type birds. Statistics were performed using SPSS24 statistics (version 24.0.0.0) software (IBM, USA).

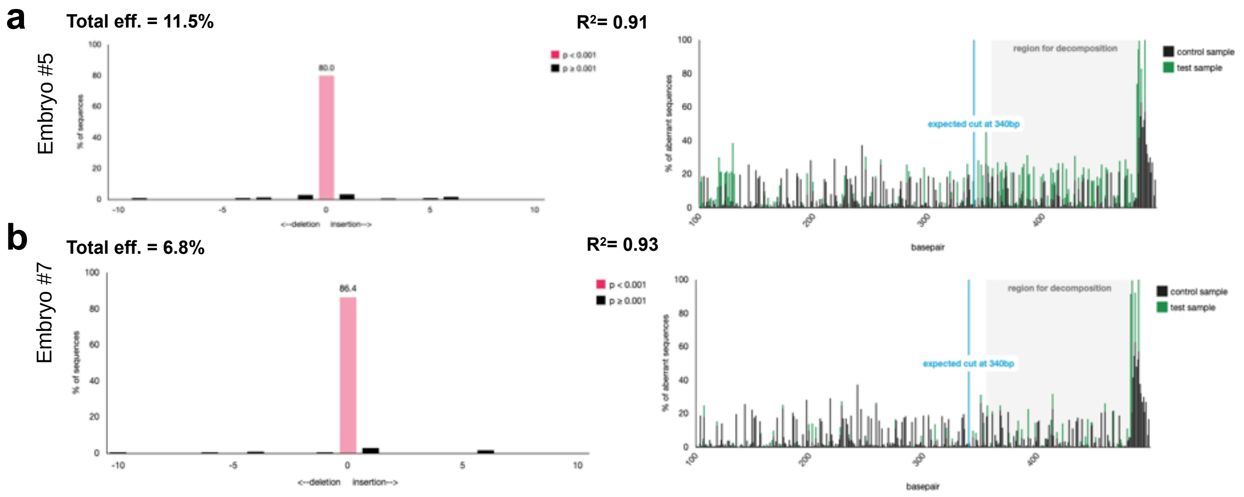


S4: TIDE analyses of *in ovo*-electroporated embryo midbrains. a, b) Analyses of the midbrain of two Cas9 positive embryos (#5, #7,) electroporated with pBlueScript II SK (+) vector containing a sgRNA against EGFP. A fragment across the EGFP target site was amplified by PCR and INDEL analysis performed.

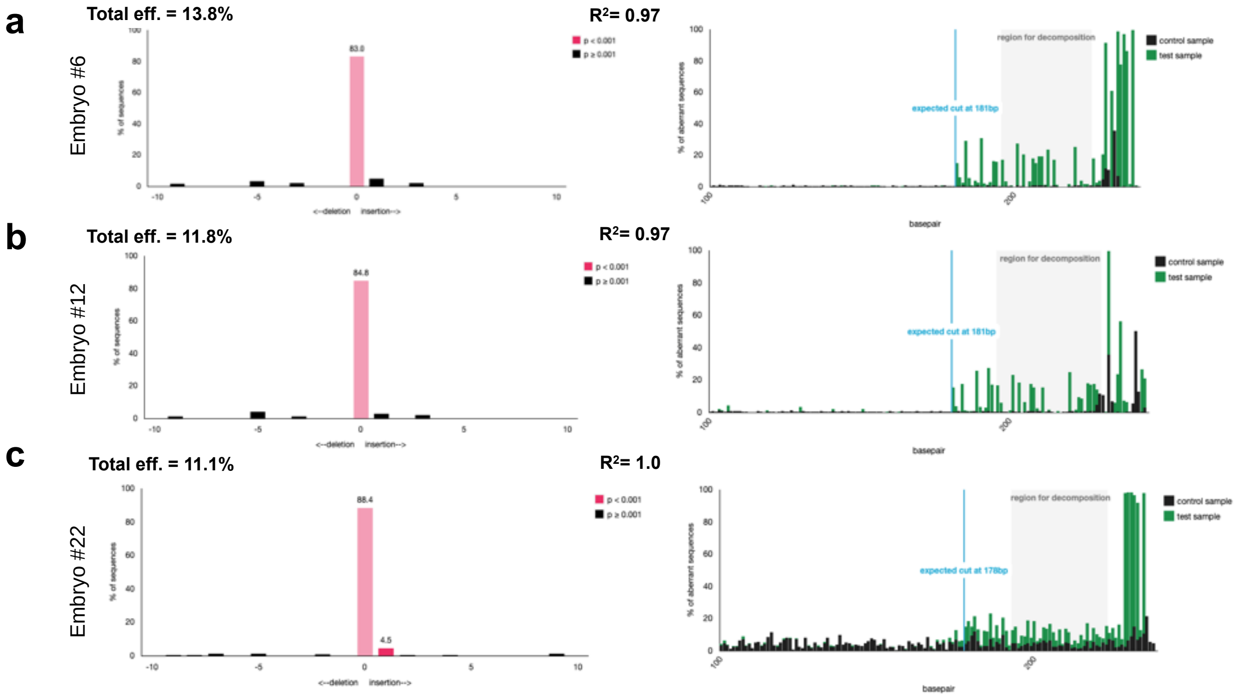


S5: TIDE analyses of *in ovo*-electroporated embryo midbrains. a, b, c) Analyses the midbrains of three Cas9 positive embryos (#6, #12, #22) electroporated with pBlueScript II SK (+) vector containing a sgRNA against *B2M*. A fragment across the B2M target site was amplified by PCR and INDEL analyses performed.
